# A novel experimental system reveals immunoregulatory responses as mediators of persistent orthohantavirus infections in a rodent reservoir host

**DOI:** 10.1101/831214

**Authors:** Tomas Strandin, Teemu Smura, Paula Ahola, Kirsi Aaltonen, Tarja Sironen, Jussi Hepojoki, Isabella Eckerle, Rainer G. Ulrich, Olli Vapalahti, Anja Kipar, Kristian M. Forbes

## Abstract

Orthohantaviruses are globally emerging zoonotic pathogens. Human infections are characterized by an overt immune response that is efficient at counteracting virus replication but can also cause severe tissue damage. In contrast, orthohantavirus infections in rodent reservoir hosts are persistent and asymptomatic. The mechanisms facilitating asymptomatic virus persistence in reservoir hosts are not well understood but could help to guide therapeutic strategies for human infections. Here we report on a study using *in vivo* and *in vitro* experiments to investigate immune responses associated with persistent Puumala orthohantavirus (PUUV) infections in the bank vole (*Myodes glareolus*), its reservoir host. We examined adaptive cellular and humoral responses by quantifying changes in T-cell related gene expression in the spleen and immunoglobulin (Ig) responses in blood, respectively. Since existing Vero E6-cell adapted hantavirus isolates have been demonstrated to have lost their wild-type infection characteristics, infections were conducted with a novel PUUV strain isolated on a bank vole cell line. Whole virus genome sequencing revealed that only minor sequence changes occurred during the isolation process, and critically, experimental infections of bank voles with the new isolate resembled natural infections. *In vitro* infection of bank vole splenocytes with the novel isolate demonstrated that PUUV promotes immunoregulatory responses by inducing interleukin-10, a cytokine strongly associated with chronic viral infections. A delayed virus-specific humoral response occurred in experimentally infected bank voles, which is likely to allow for initial virus replication and the establishment of persistent infections. These results suggest that host immunoregulation facilitates persistent orthohantavirus infections in reservoir hosts.

**Importance:** Orthohantaviruses are a group of global pathogens that regularly spillover from rodent reservoirs into humans and can cause severe disease. Conversely, infections in reservoir hosts do not cause obvious adverse effects. The mechanisms responsible for persistent asymptomatic reservoir infections are unknown, and progress has been hindered by the absence of an adequate experimental system. Knowledge on these mechanisms could help provide strategies to treat human infections. We developed and validated an experimental system based on an orthohantavirus isolated in cells of its vole reservoir host. Using animal and cell culture experiments in the reservoir host system, we demonstrated that infection suppresses immunity in the vole reservoir via specific mechanisms, likely allowing the virus to take hold and preventing immune responses that can cause self-damage.

## Introduction

Understanding how zoonotic pathogens are maintained and transmitted in nature is critical for efforts to curtail human disease (1). Most investigations have focused on the identification of wildlife reservoir hosts (2, 3), and to a lesser extent, the characterization of high-risk subgroups and time periods, such as migration and breeding periods (4, 5), when transmission is greatest in reservoir host populations and the risk of human exposure may therefore be elevated. However, it remains largely unknown how zoonotic pathogens are maintained at an individual-level in wildlife reservoirs, such as the molecular and cellular mechanisms of host-pathogen interactions that promote clearance or persistence, and feed back onto these population-level dynamics.

Hantaviruses (genus *Orthohantavirus*, order *Bunyavirales*) provide one of the most clearly described reservoir host-zoonotic pathogen relationships (6). Hantavirus species have been identified across the globe, some of which cause Hemorrhagic Fever with Renal Syndrome (HFRS) in Eurasia or Hantavirus Cardiopulmonary Syndrome (HCPS) in the Americas (7). These diseases begin abruptly with a general febrile phase followed by signs of vascular leakage, predominantly manifesting as transient kidney failure in HFRS or as life-threatening pulmonary complications in HCPS (7, 8). The pathogenesis of both conditions is immune-mediated, with pronounced activation of immune cells and pro-inflammatory cytokines during acute stages of disease. An intense hantavirus-directed immune response then results in efficient virus clearance in surviving patients (8).

In contrast to human infection, hantavirus infections are thought to be persistent and asymptomatic in their wildlife reservoirs (6). Each hantavirus species is carried by a specific reservoir host; for example, Puumala orthohantavirus (PUUV), which is present in Europe and causes thousands of human infections annually (9), is carried by the bank vole (*Myodes glareolus*). Based on mitochondrial cytochrome *b* gene sequences different evolutionary lineages of the bank vole with different glacial refugia were defined (10–13). In southern to central Finland the Eastern evolutionary lineage has been identified (13).

Persistent infections are generally facilitated by host immune tolerance (14), and it has been suggested that activation of regulatory T cells in reservoir hosts could also play a role in enabling hantavirus persistence (15, 16). Deciphering the detailed molecular mechanisms by which reservoir hosts become immune tolerant instead of attempting potentially costly virus clearance could significantly increase our understanding of hantavirus pathogenesis and provide insights into alternative treatment strategies.

A major issue hindering detailed research on the molecular mechanisms of hantavirus persistence in reservoir hosts has been the absence of a standardized experimental infection system. Two main experimental strategies have been employed by researchers to date: 1) lung homogenates from naturally infected rodent hosts have been used as virus inoculum. However, this method suffers from slowly developing infections and a lack of standardized infection loads (17, 18), and 2) hantaviruses isolated and grown to high titers in cell culture. However, this method has been performed so far almost exclusively using Vero E6 cells, which generates cell culture-adapted viruses with attenuated properties (19–22).

The purposes of this study are to develop an effective hantavirus experimental wildlife system and to examine the immunologic responses that facilitate persistent hantavirus infections in a reservoir host species. To these means we employed PUUV and its reservoir host, the bank vole. Our first goal was to isolate PUUV from wild bank voles using a host-derived cell line, and to evaluate genotypic changes during the isolation process using next-generation sequencing (NGS). Experimental bank vole infections were then used to compare the viral RNA load and tissue distribution and) of the novel isolate to the wild bank vole-derived PUUV–containing lung homogenate (PUUV-wt), a Vero E6-cell attenuated PUUV-Kazan strain and to PUUV in naturally infected wild voles.

To examine immunologic responses to PUUV in vole reservoir hosts, we measured T helper cell differentiation pathways by assaying changes in mRNA expression of cytokines and transcription factors (TF) related to T helper (Th) 1-, Th2- and regulatory T (Treg) cell activation. Comparisons were made among voles infected with the novel PUUV isolate, PUUV-Kazan, PUUV-wt and mock infected voles. Since no assays to detect PUUV-specific T cell responses are available for bank voles, the bulk expression levels of T cell-related mRNAs in the spleen were studied both directly in spleens of experimentally infected voles and using single-cell suspensions (splenocytes) from PUUV-negative bank voles after *in vitro* infection with PUUV isolates. To gain further insights into PUUV-specific and non-specific humoral responses to PUUV infections, we additionally measured PUUV nucleocapsid (N)-specific and total immunoglobulin G (IgG) levels in vole blood following infection.

## Results

### Isolation of PUUV-Suo in epithelial bank vole cells

A renal epithelial cell line Mygla.REC.B of the Western evolutionary lineage of the bank vole was generated to isolate a new PUUV strain named Suonenjoki (PUUV-Suo) from the lungs of a PUUV-infected wild bank vole collected in Suonenjoki, Finland. Based on the detection of PUUV N protein by immunofluorescence asssay (Fig. 1), 100% of cells were infected after approximately 20 cell passages post initial inoculation with the PUUV-containing lung homogenate. Cell passaging did not influence virus infectivity, as confirmed by consistent detection of progeny viruses in cell culture supernatants at 10^4^ fluorescence focus forming units (FFFUs)/ml.

**Figure 1.**
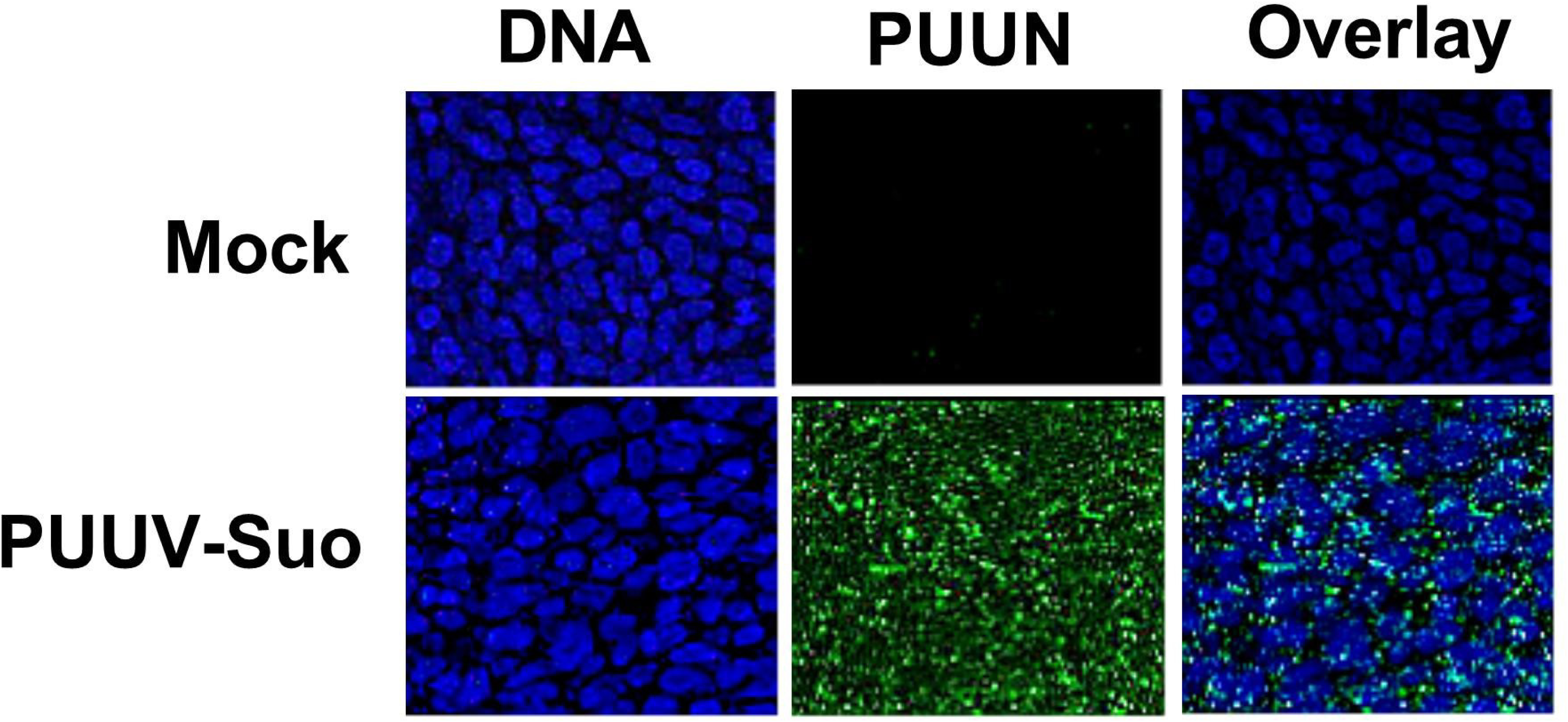
Immunofluorescence image showing successful isolation of PUUV-Suo in a bank vole cell line. Non-infected (mock) or PUUV-Suo infected bank vole kidney epithelial cells (Mygla.REC.B) were stained for nuclei (DNA) using Hoechst 33420 (blue) and for PUUV nucleocapsid protein (PUUN) using PUUN-specific rabbit polyclonal antibody followed by AlexaFluor488-conjugated secondary antibody (green).

The isolation process resulted in eight nucleotide exchanges when compared to the consensus sequence derived from the lung homogenate (Table 1). Two mutations occurred in non-coding regions, three resulted in amino acid exchanges in the RNA-dependent RNA polymerase (RdRP, L segment), one resulted in an amino acid exchange in the N protein, and two were silent mutations in the glycoprotein precursor (GPC; M segment) and RdRP coding regions. The new virus isolate had a 95-99 % nucleotide sequence identity with partially sequenced PUUV strains from bank voles in Konnevesi and from a fatal human case in Pieksämäki (Supplementary Figs 1-3) (23, 24), both located within 100 km of the vole trapping site in central Finland.

**Table 1.** Comparison between the consensus nucleotide sequences and amino acid sequences of the PUUV-Suo strain at the pre-isolation stage in the lungs of a naturally infected bank vole and after isolation in Mygla.REC.B cells.

| Segment | Nucleotide position in anti-genomic orientation | Nucleotide in original virus | Nucleotide in isolate | Amino acid in original virus | Amino acid in isolate | Codon position |
| --- | --- | --- | --- | --- | --- | --- |
| L | 66 | A | G | no change |  | 3rd |
|  | 411 | G | A | Methionine | Isoleucine | 3rd |
|  | 5700 | A | G | Isoleucine | Methionine | 3rd |
|  | 5942 | C | U | Alanine | Valine | 2nd |
| M | 89 | U | C | no change |  | 1st |
|  | 3544 | U | C |  |  | non-coding |
| S | 136 | G | U | Valine | Leucine | 1st |
|  | 1625 | C | U |  |  | non-coding |

### Replication of PUUV-Suo in *Mygla.REC.B cells* is resistant to immune stimulation

We hypothesized that the observed persistence of PUUV-Suo in bank vole cells would be a consequence of either impaired host cell antiviral signaling or the ability of the virus to counteract antiviral responses. Therefore, the antiviral function of Mygla.REC.B cells was assessed by relative reverse transcription-real time quantitative polymerase chain reaction (RT-qPCR) quantification of the interferon-inducible myxovirus resistance protein 2 (Mx2) mRNA after exogenous activation of non-infected and infected cells with the toll-like receptor 3 (TLR3) agonist polyI:C or with Sendai virus (SeV, also known as Murine respirovirus). Firstly, supporting the immune competence of Mygla.REC.B cells, a significant upregulation of Mx2 mRNA occurred in non-infected cells with both stimulants (~100-fold and 2000-fold by polyI:C and Sendai virus, respectively, Fig. 2A). Secondly, non- and polyI:C-stimulated PUUV-infected cells showed higher levels of Mx2 mRNA than non-infected cells. PUUV and Sendai virus superinfection had a significant synergistic effect on Mx2 mRNA production (Fig. 2A). These findings suggest that Mygla.REC.B cells are capable of interferon signaling but that persistent PUUV-Suo infection does not impair this pathway.

**Figure 2.**
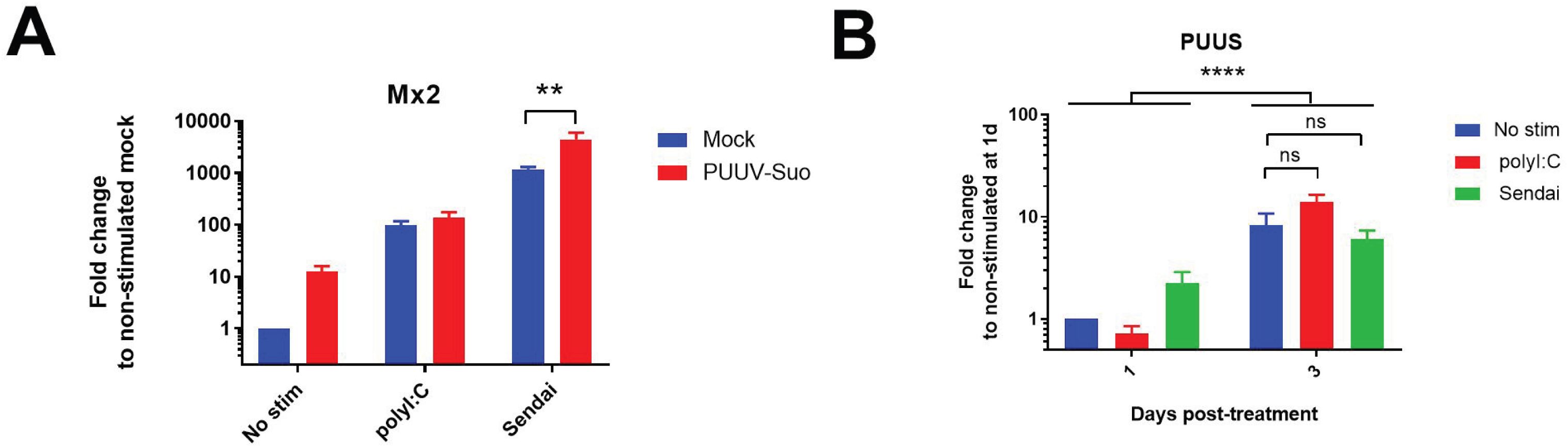
Interaction between PUUV-Suo infection and innate immune pathway stimulation in Mygla.REC.B cells. (**A**) Non-infected (mock) or PUUV-Suo infected Mygla.REC.B cells were either non-stimulated (No stim), stimulated with polyI:C or infected with Sendai virus (Sendai) for 1 day. RNA was isolated from cells and subjected to relative quantification of Mx2 mRNA using RT-qPCR. (**B**) PUUV-Suo infected Mygla.REC.B cells were either non-stimulated, stimulated with polyI:C or infected with Sendai virus for 1 and 3 days. RNA was isolated from cells and subjected to relative quantification of PUUV S segment RNA using RT-qPCR. ** (p < 0.01) or **** (p < 0.001). ns, not significant. Bars represent mean +/- standard deviation (n = 2).

We also investigated whether PUUV replication in Mygla.REC.B cells is susceptible to an exogenously induced antiviral state. PUUV-infected cells, non-stimulated or stimulated with polyI:C or Sendai virus, were assessed for PUUV S segment RNA levels by RT-qPCR at 1 and 3 days (d) post-stimulation. Neither polyI:C nor Sendai virus stimulation influenced PUUV S segment RNA levels, which increased significantly from days 1 to 3 with and without stimulation (Fig. 2B), indicating that PUUV-Suo infection in Mygla.REC.B cells is not affected by innate immune pathway stimulation.

### Experimental PUUV-Suo infection resembles natural infection in bank voles

Following experimental infection of bank voles with PUUV-Suo or a mock inoculum (PBS), the amount of PUUV S segment RNA was measured in lungs, spleen, kidney and blood of voles at 3d, and 1, 2 and 5 weeks (wks) post infection (pi). Viral RNA was similarly measured in the urine of experimentally infected voles at 3d, and 1 and 2wks pi. As another control, virus RNA load and RNA distribution in tissues were also assessed at 2 and 5wks pi from voles inoculated with UV-inactivated PUUV-Suo isolate.

PUUV RNA was detected in all organs of PUUV-Suo infected voles from 3 dpi onwards (Fig. 3A), but not in mock-infected animals nor those inoculated with UV-inactivated virus (data not shown). Highest PUUV RNA levels occurred in the lungs at 3d and 1wk pi and decreased thereafter, with increased levels then observed in the spleen (at 2 and 5wk pi) and kidneys (at 5wk pi). PUUV RNA was detected in the blood of one vole at 3 dpi, suggesting that if PUUV-Suo caused viremia it occurred very early during infection. We also detected PUUV RNA in the urine of all tested PUUV-Suo infected bank voles (n = 7), suggesting virus shedding.

**Figure 3.**
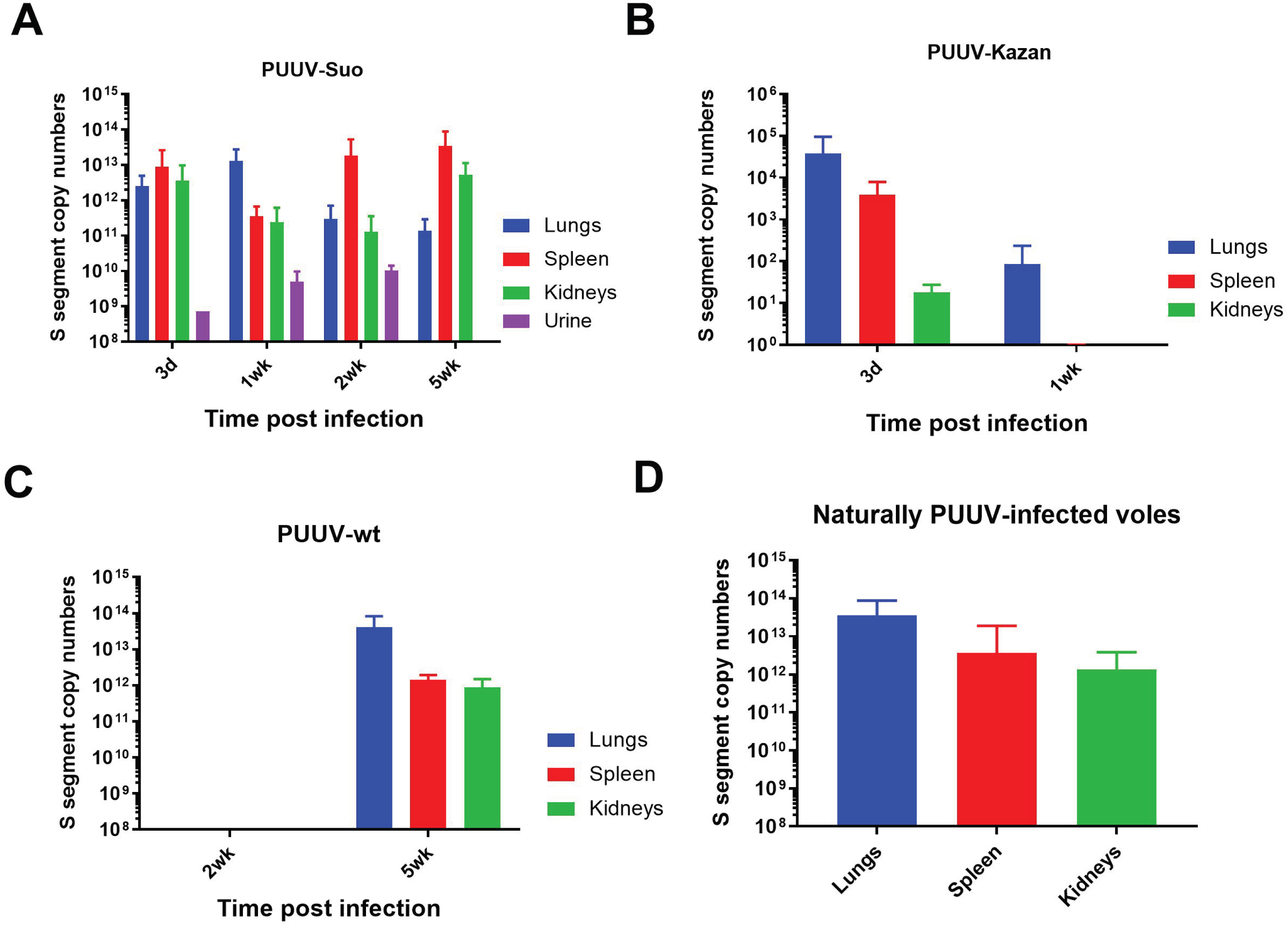
Viral RNA load analysis of experimentally and naturally infected bank voles. The lungs, spleen and kidneys of bank voles infected with PUUV-Suo (**A,** n = 4), PUUV-Kazan (**B,** n = 3) or PUUV-wt (**C,** n = 3) were collected at indicated times post infection to quantify PUUV copy numbers by PUUV S segment-specific qPCR. In addition, seropositive naturally PUUV-infected bank voles with unknown infection timepoints were included as reference (**D,** n = 27-38). Bars represent mean + standard deviation.

We further compared the infection dynamics of the PUUV-Suo isolate to a Vero E6 cell-adapted PUUV-Kazan isolate, PUUV-wt and to natural PUUV infections. Low levels of PUUV RNA were detected following infection with PUUV-Kazan, mainly in the lungs of infected voles at 3d pi; after this the virus was efficiently cleared, with viral RNA levels significantly decreased at 1wk pi (Fig. 3B). No virus replication was observed for PUUV-wt at 2wk pi, but three out of four voles were positive at 5wk pi (Fig. 3C). At this time point, viral RNA loads were highest in the lungs, followed by the spleen and kidneys. The amount and distribution of PUUV RNA seen in naturally PUUV-infected wild bank voles (with unknown infection timepoint) resembled that seen in PUUV-Suo infected voles during the first week and in voles inoculated with PUUV-wt at 5wk pi (Fig. 3D).

### PUUV-Suo host cell range resembles natural infections

Both naturally and experimentally infected bank voles were examined by histology and immunohistology for PUUV N protein. The two naturally infected voles did not exhibit any histopathological changes and viral antigen expression was limited in its amount and distribution. Besides endothelial cells in renal glomerula and in some capillaries of the liver, kidney, lungs and heart (Fig. 4A, B), pneumocytes (mainly type II) and macrophages in liver (Kupffer cells) and spleen (red pulp macrophages) were found to be occasionally positive (Fig. 4B). In one animal with a higher number of positive cells, viral antigen was also detected in tubular epithelial cells in the renal cortex and medulla.

**Figure 4.**
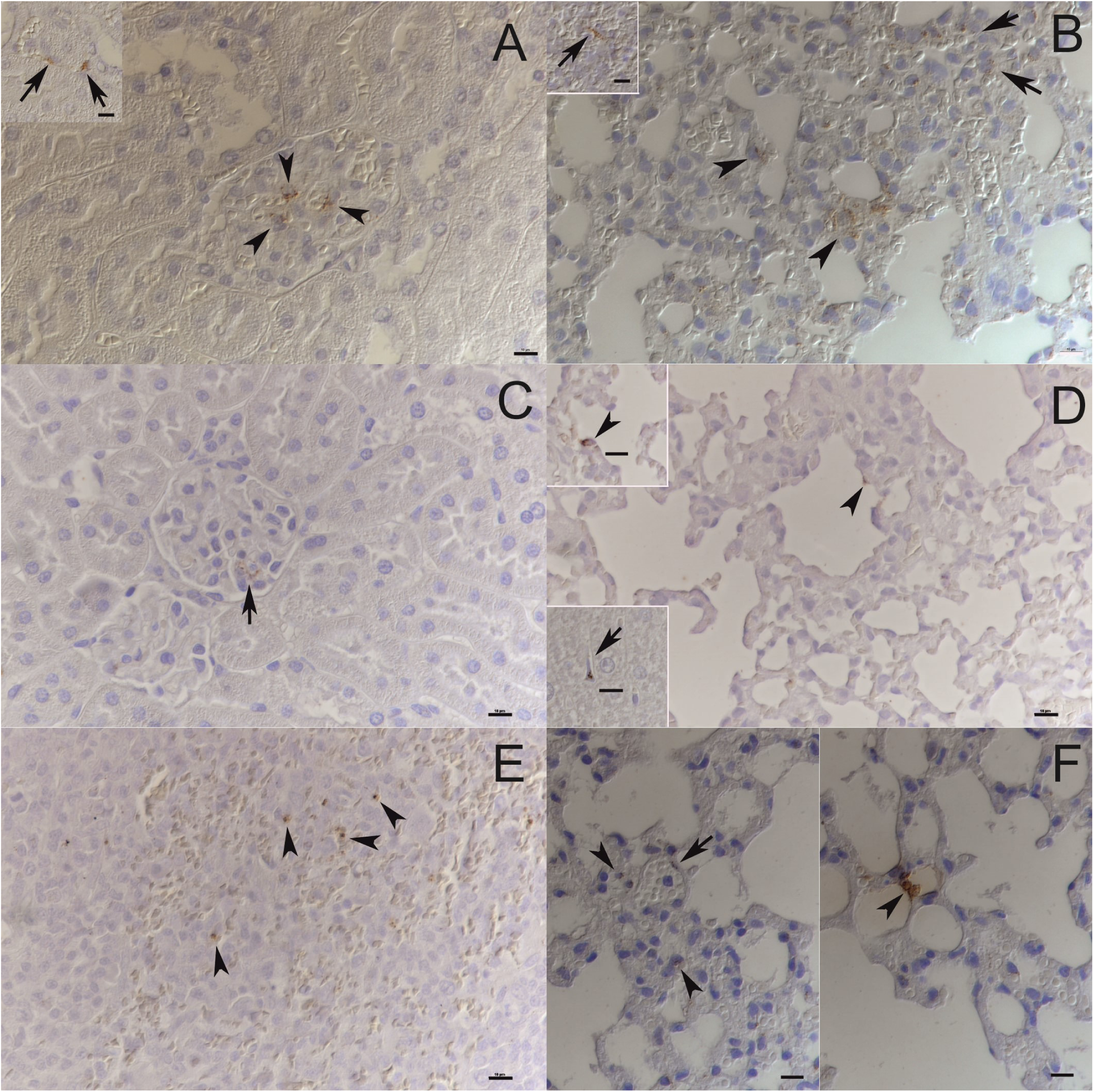
PUUV target cell analysis in experimental and natural infections. (**A)** Kidneys of naturally infected bank voles: Viral antigen expression is apparent in occasional glomerular endothelial cells (arrowheads) and occasional vascular endothelial cells (inset: arrows). (**B)** Lungs and spleen of naturally infected bank voles: A variable number of vascular/capillary endothelial cells (arrows) and pneumocytes (arrowheads) express viral antigen in the lungs. Very few cells in the splenic red pulp, consistent with macrophages, are also found to express viral antigen (inset: arrow). (**C)** Kidney of PUUV-Suo infected bank vole at 1 wk pi show scattered glomerular endothelial cells with viral antigen expression. (**D)** PUUV-Suo infected vole at 3 d exhibit occasional positive alveolar epithelial cells in the lungs (arrowhead, also in top inset), and scattered positive Kupffer cells in the liver (bottom inset: arrow). (**E)** The spleen of PUUV-Suo infected voles at 2 wks pi show very limited viral antigen expression, which is restricted to individual macrophages, for example in the splenic red pulp (arrowheads). (**F)** The lungs of PUUV-wt infected bank vole at 5 wks pi show viral antigen expression restricted to occasional vascular endothelial cells (left, arrow) and scattered pneumocytes in the lungs (right, arrowhead). The images are representative of two voles analyzed by immunohistochemistry for PUUV N protein expression in naturally infected voles and at each time point of PUUV-Suo or PUUV-wt infections. Haematoxylin counterstain. Bars = 10 µm.

None of the experimentally PUUV-Suo infected voles exhibited histopathological changes (two voles analyzed at each time point). During the early stages of infection (≤ 1wk pi), the infection pattern was similar to that seen in naturally infected bank voles. Capillary endothelial cells (also in glomerular tufts), pneumocytes and macrophages (Kupffer cells in the liver, pulmonary alveolar macrophages) occasionally exhibited viral antigen expression (Fig. 4C, D). At later time points (2 and 5wks pi), viral antigen was only found in macrophages, and almost exclusively in rare macrophages of the splenic red pulp (Fig 4E). No PUUV N protein was detected in PUUV-Kazan-infected voles, consistent with the low level of PUUV RNA in those animals (Fig. 3C).

Histological examination of the PUUV-wt infected voles at 5wks pi did not reveal any pathological changes. Interestingly, when testing lungs, kidneys and spleen by immunohistochemistry, N protein expression was only detected in one of three animals, and only in the lungs, where a few individual pneumocytes and endothelial cells were positive (Fig. 4F). This animal also showed the highest viral RNA levels in the lungs.

### PUUV-specific immunoglobulin (Ig) responses are delayed during persistent infections

All PUUV-Suo and PUUV-Kazan infected voles, which were infected for at least one week, seroconverted to produce PUUV-specific Ig but with contrasting kinetics (Fig. 5A; the assay does not discriminate between Ig-subclasses). For PUUV-Suo, specific Ig-titers were low at 1wk but peaked at 2wks pi. This was significantly different from PUUV-Kazan infected voles, in which highest PUUV-specific Ig titers were detected at 1wk pi. For PUUV-wt, only one of three infected voles had seroconverted at 3 wks pi (Fig. 5A).

**Figure 5.**
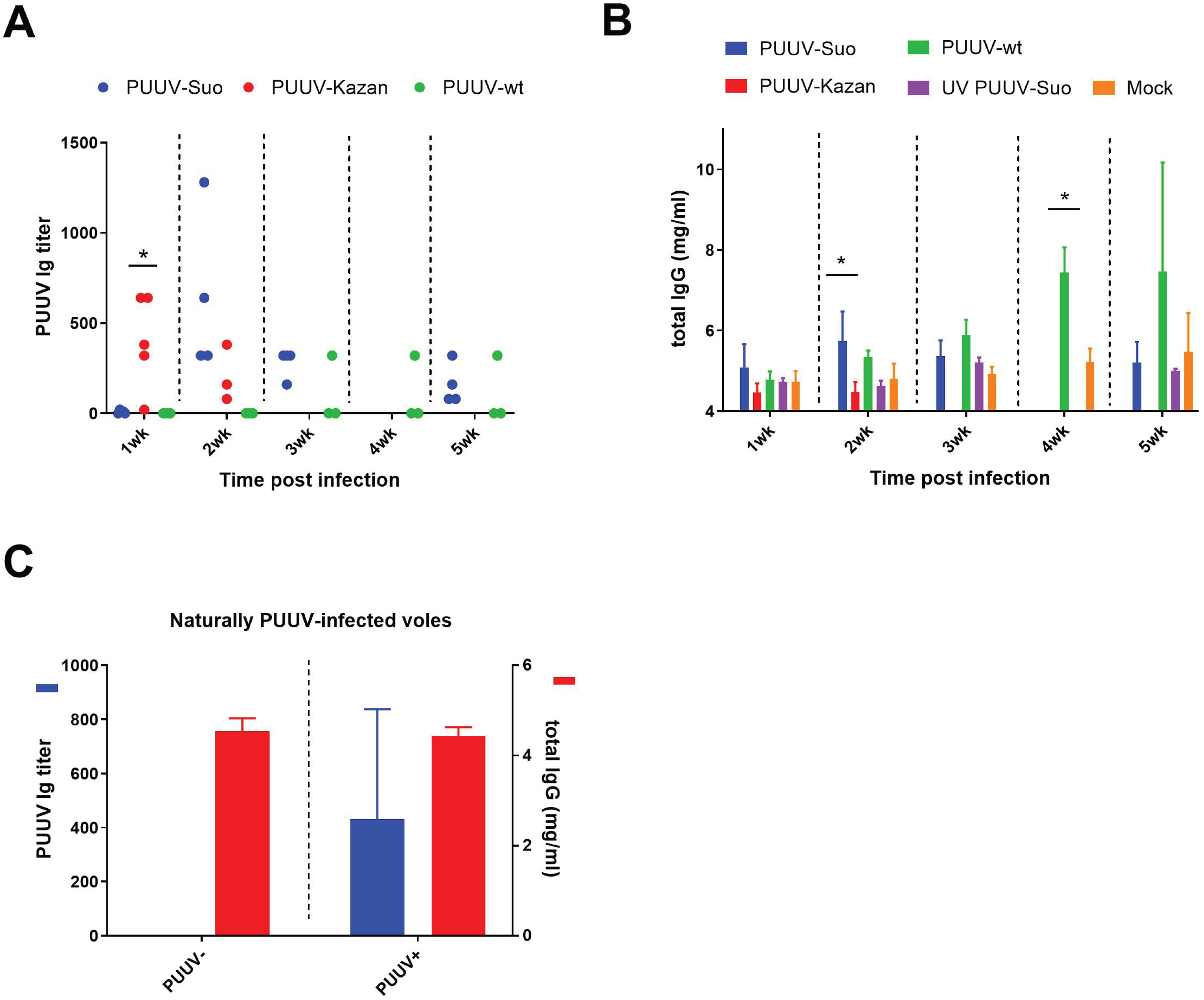
Assessment of PUUV-specific Ig and total IgG responses in blood of experimentally and naturally infected bank voles. (**A**) PUUV-specific Ig titers in blood of PUUV-Suo (n = 4), PUUV-Kazan (n = 3-5) and PUUV-wt (n = 3) infected voles at indicated time points. (**B**) Total IgG levels in blood of PUUV-Suo (n = 4-12), PUUV-Kazan (n = 2) and PUUV-wt (n = 2) infected voles at indicated time points. As a comparison, blood of mock-(n = 2-8) and UV-inactivated PUUV-Suo (n = 2) infected voles were also assessed at the indicated time points. (**C**) PUUV-specific Ig titers and total IgG levels in blood of naturally PUUV-infected or non-infected bank voles with unknown infection timepoints (n = 8). Bars represent mean +/- standard deviation. * indicates statistically significant difference (p < 0.05).

Next we measured total IgG levels in the blood of experimentally infected voles. Compared to mock-infected animals at 2wks pi, voles infected with PUUV-Suo showed elevated total IgG levels, while no difference was observed between voles infected with the PUUV-Kazan or the mock inoculate at 1wk and 2wks pi (Fig. 5B). The two PUUV-wt infected voles analyzed had elevated total IgG levels when compared to mock-infected voles (Fig. 5B), despite only one showing a PUUV-specific response (Fig. 5A).

To gain further insights into whether total IgG levels were comprised predominantly of PUUV-specific responses in experimentally infected voles, PUUV-specific Ig and total IgG titers were assessed also in naturally PUUV-infected voles. Despite detecting PUUV-specific responses at levels comparable to experimentally PUUV-Suo infected voles, total IgG in naturally infected voles was not elevated when compared to non-infected voles (Fig. 5C), suggesting that the acute total IgG response towards PUUV-Suo infection involves PUUV non-specific IgG.

### Minor effects of PUUV infection on splenic T cell related gene expression

To assess the potential effects of experimental PUUV infection on T cell differentiation, we assayed pre-selected splenic T cell related mRNAs in mock-, PUUV-Suo and PUUV-wt infected voles for transcriptional regulation (the same animals as those examined by PUUV-specific RT-PCR in Fig. 3). Transcription levels of cytokines associated with Th1, Th2 or Treg cell activation (interferon {IFN}-γ, interleukin {IL}-4, IL-10 or transforming growth factor {TGF}-β) did not differ significantly between PUUV- and mock-infected voles (Fig. 6). Of key transcription factors for Th1, Th2 and Treg pathways (Tbet, Gata binding protein 3 {Gata3} or Forkhead box P3 {FoxP3}, respectively), FoxP3 expression was significantly lower in PUUV-wt infected than in mock-infected voles at 5 wks dpi. A similar trend, although not statistically significant, was observed for PUUV-Suo at 2 and 5 wks dpi. Another significant change in immune gene regulation was represented by the upregulation of Mx2 gene expression in response to PUUV-Suo infection in comparison to mock infection at 5wks pi (Fig. 6B). A trend towards higher Mx2 mRNA levels in response to PUUV-wt was also observed at the same time point (Fig. 6C).

**Figure 6.**
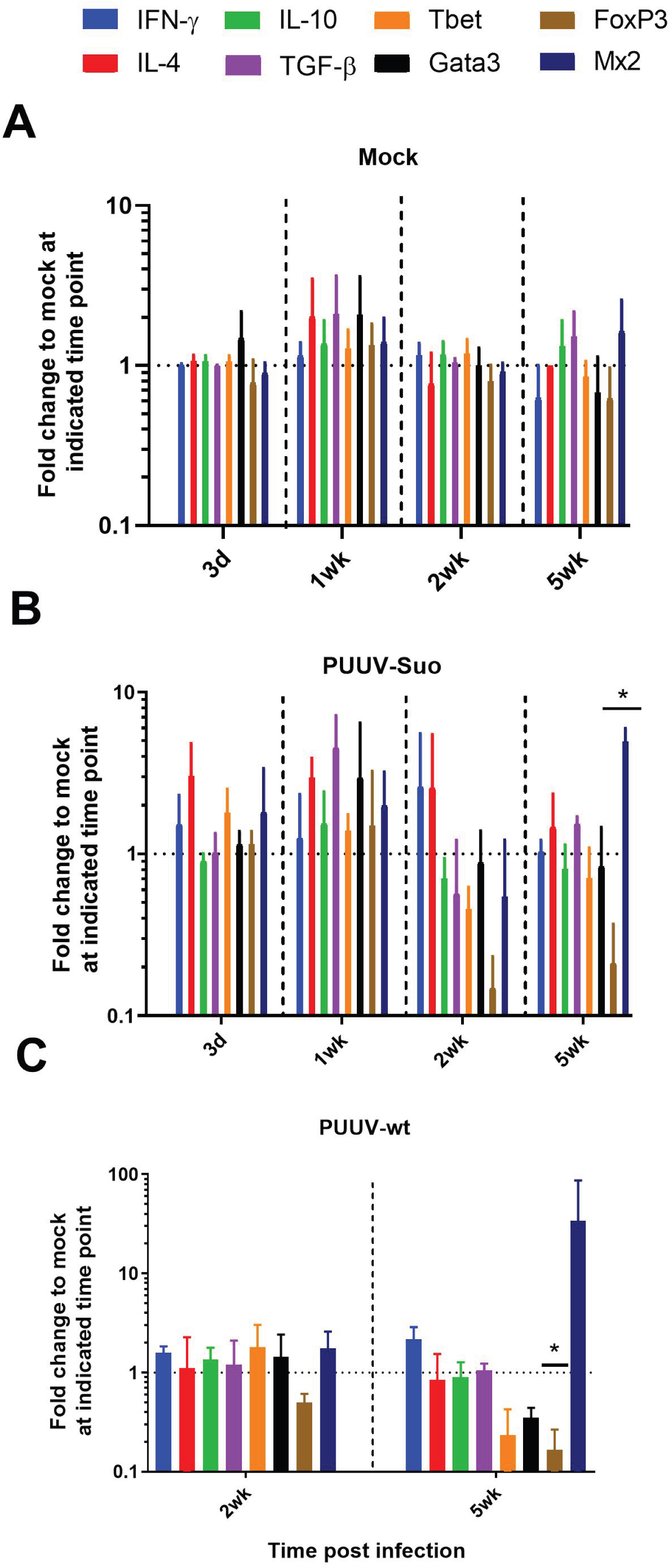
Comparison of T-cell related transcription factor and cytokine mRNA levels in experimentally PUUV-infected voles. The spleens of mock (**A,** n = 2), PUUV-Suo (**B,** n = 4), and PUUV-wt (**C,** n = 3) infected voles were subjected to relative mRNA expression level analysis by RT-qPCR at indicated time points. Statistically significant differences were assessed separately for each gene between mock- and PUUV-infected voles. * (p < 0.05). Bars represent mean +/- standard deviation.

### PUUV-Suo promotes immunoregulation via cytokine IL-10

Splenocytes isolated from PUUV-negative wild-caught bank voles were inoculated *in vitro* with PUUV-Suo, PUUV-Kazan and UV-inactivated PUUV-Suo and assessed for changes in T cell-related gene expression. IFN-γ mRNA levels were higher in PUUV-Kazan infected splenocytes than in PUUV-Suo infected splenocytes (~11-fold vs. ~3-fold increase in comparison to non-infected controls, respectively; Fig. 7). In contrast, IL-10 mRNA expression was significantly upregulated in response to PUUV-Suo when compared to PUUV-Kazan and UV-inactivated PUUV-Suo (~6-fold vs. 2-3-fold increase in comparison to non-infected controls, respectively). No other significant differences in transcription levels of other genes (TGF-β, Mx2, T-bet, Gata3) were observed. However, we were unable to measure IL-4 and FoxP3 mRNA, likely due to the generally lower RNA levels in single-cell suspensions than in tissue samples. The PUUV-specific RT-PCR confirmed that live viruses (PUUV-Suo and Kazan), but not UV-inactivated viruses replicated in bank vole splenocytes at a comparable level; however the level of infection varied significantly among splenocyte donors (data not shown).

**Figure 7.**
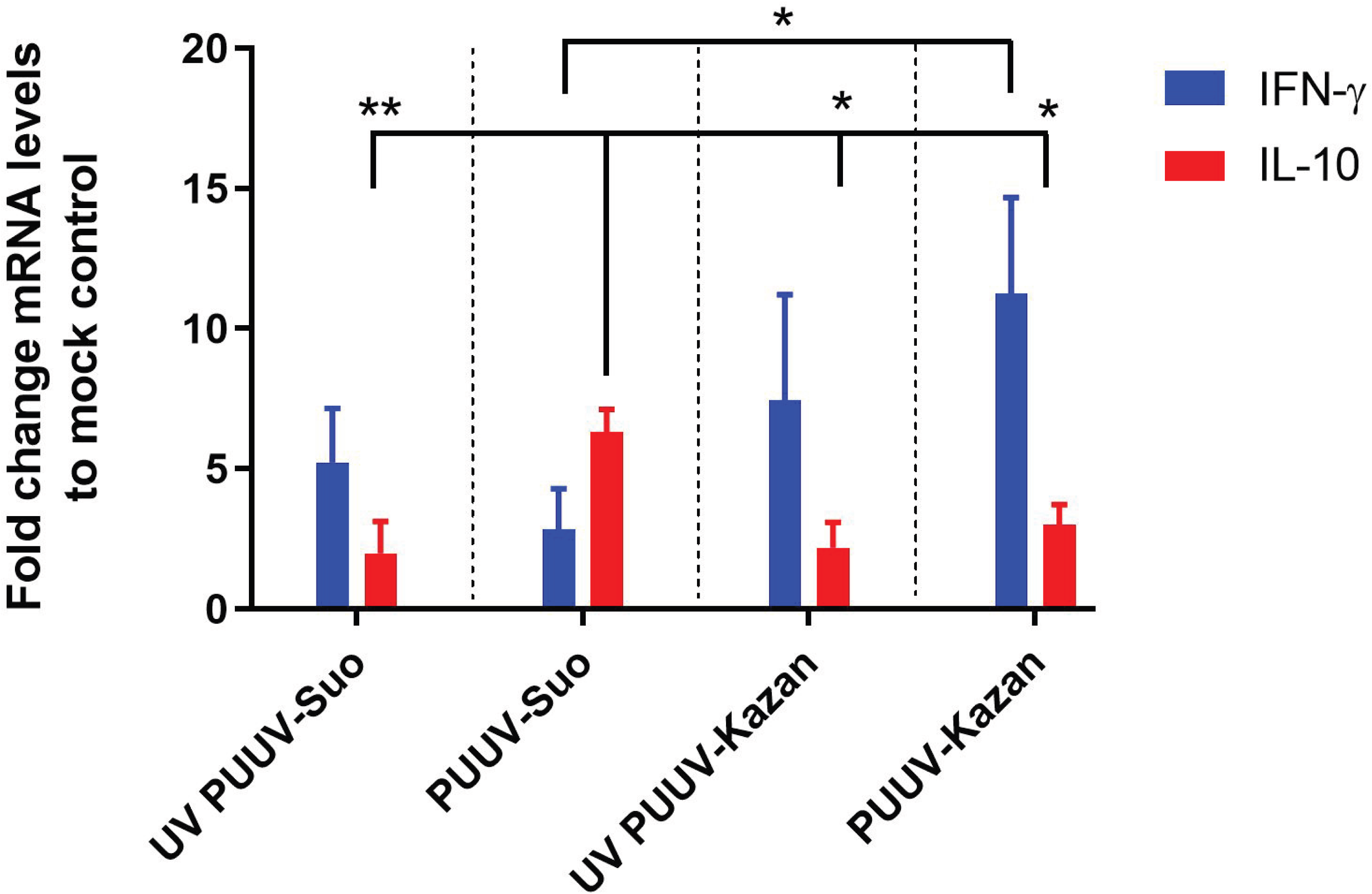
Comparison of cytokine mRNA responses in bank vole splenocytes infected with PUUV-Suo and PUUV-Kazan isolates. Splenocytes were isolated from PUUV-negative bank voles and either infected with UV-inactivated or live PUUV isolates or not infected (n = 3 for PUUV-Suo; n = 2 for PUUV-Kazan; multiplicity of infection 0.01 for both). Quantification of mRNAs was conducted relative to non-infected splenocytes using RT-qPCR. * (p < 0.05) or ** (p < 0.01). Bars represent mean +/- standard deviation.

## Discussion

In this study we established and validated an experimental hantavirus rodent reservoir model. Using a virus newly isolated on a bank vole cell line, we conducted standardized experimental *in vivo* and *in vitro* infections to examine immune responses at cellular and organismic levels, gaining insight into the mechanisms of PUUV persistence in the reservoir host. These experiments revealed that activation of the immunoregulatory cytokine IL-10 differentiates wild-type PUUV from attenuated isolates, which is likely to be a key factor in hantavirus persistence. Experimentally infected voles also exhibited a delayed PUUV-specific humoral response, suggesting that PUUV has developed mechanisms to evade host immune recognition.

The isolation of hantaviruses in cell culture is rare and has so far typically been achieved using interferon type 1-deficient cells (Vero E6 cells) derived from African green monkeys (25), which result in attenuated wild-type properties of the virus (19–22). The PUUV-Suo strain is the first hantavirus isolated on a cell line of the respective natural reservoir host. The isolation process of PUUV-Suo in bank vole renal epithelial cells resulted in a significantly lower nucleotide substitution frequency (0.06 %) when compared to a PUUV-Kazan strain on Vero E6 cells (21, 26) and persistent cell culture infections that were refractory to exogenous stimulation of innate immune pathways. Importantly, the PUUV-Suo isolate caused persistent vole infections without obvious pathological effects. Target cell patterns and viral RNA loads, particularly during the early stages of infection, were consistent with those seen in naturally infected voles and were considerably different from experimental infections conducted with the attenuated PUUV-Kazan isolate. The MyGla.REC.B cell line used to isolate PUUV-Suo is derived from the Western evolutionary lineage, which is different from the Eastern bank vole lineage present in central Finland (27) and used as the source and target of PUUV-Suo isolate. The efficient replication of the Eastern bank vole lineage-derived PUUV-Suo strain in Western lineage bank vole-derived cell line is in line with results from a field study in Germany demonstrating transmission of Western lineage-associated PUUV to sympatrically occurring bank voles of the Eastern and Carpathian lineages (28). Further in vitro studies are needed to determine whether heterologous bank voles evolutionary lineages could support PUUV isolation and/or infection. These future investigations may profit from the recent establishment of permanent bank vole cell lines from different evolutionary lineages of the bank vole (29).

The viral target cell pattern observed for PUUV-Suo is similar to, but less widespread than in weanling and suckling bank voles that were inoculated with lung homogenates from PUUV-positive bank voles (17). Experimental infections conducted with Sin Nombre orthohantavirus (SNV) in deer mice yielded comparable, though apparently more intense viral antigen expression (18). In naturally infected deer mice, however, only pulmonary and cardiac vascular endothelial cells were found to be positive, with an expression intensity that resembled that observed in the present study (30). Interestingly, only PUUV-Suo appears to infect renal tubular epithelial cells, which would account for more intense virus shedding in urine. This could reflect different primary transmission routes for PUUV and SNV, with PUUV suggested to be shed more in urine and SNV in the saliva (6).

In support of the ability of PUUV to evade T cell immune responses, we detected no significant upregulation in mRNA expression (including transcription factors Tbet, Gata3, FoxP3 and cytokines IFN-γ, IL-4, IL-10 and TGF-β) in the spleens of bank voles infected with PUUV-Suo or wild PUUV-containing lung homogenates (PUUV-wt). Instead, a decrease in FoxP3 levels in PUUV-wt infected voles is contrary to the suggested role of the Tregs in mediating virus persistence, as detected in other experimental hantavirus-reservoir host interactions (31, 32). These discrepant results could be explained by differences in experimental setups. While we measured bulk FoxP3 mRNA levels in the spleen, others have looked either at splenic antigen-specific (31) or lung-resident (32) Treg responses. It is possible that the decrease in FoxP3 mRNA and associated Treg levels in the spleen reflects the recruitment of these cells into peripheral sites of infection in our experimental setup. However, inherent differences in the persistence mechanisms among PUUV and other hantaviruses cannot be ruled out and needs to be further evaluated.

An interesting finding was that PUUV-Suo induced IL-10 mRNA expression in bank vole splenocyte cultures. This was not observed with attenuated PUUV-Kazan, which instead induced IFN-γ. IL-10 is well-known for its ability to hinder pro-inflammatory responses (33) and could explain the lack of IFN-γ induction in PUUV-Suo infected splenocytes. The fact that we did not observe elevated IL-10 mRNA levels in the spleen of PUUV-Suo (or PUUV-wt) infected voles could be explained by a higher multiplicity of infection (m.o.i.) achieved in an *in vitro* environment using cultured splenocytes. It is also possible that a potential increase in IL-10 transcription does occur in the spleen of infected voles, but is short-lived and thus difficult to observe using samples collected with several day intervals.

The increase in IL-10 expression due to PUUV in bank vole splenocytes is not surprising given the established role of this cytokine in promoting chronic infections (34, 35). Lymphocytic choriomeningitis virus (LCMV) infection in mice is the best-studied non-human model for persistent virus infections. For this mammarenavirus, IL-10 blocking experiments have been shown to lead to virus clearance, demonstrating the key role of IL-10 in driving persistent infections (36–38). The immunomodulatory effects of IL-10 are exerted through antigen presenting cells (APCs) which, in response to IL-10, inhibit effector Th1 cell function and downregulate inflammation (33). Our results demonstrating early induction of IL-10 in PUUV-infected splenocytes together with findings of virus-infected macrophages *in vivo* indicate a similar scenario in PUUV-infected bank voles; virus reverts early macrophage responses from a typical inflammatory milieu towards immunomodulation with subsequent inhibition of IFN-γ expression, Th1 cell activation and virus clearance. In fact, corroborating the differential response between human and reservoir host APCs to hantavirus infection, human dendritic cells have been shown to decrease IL-10 expression while elevating pro-inflammatory responses when in contact with Andes orthohantavirus-infected endothelial cells (39).

Inadequate humoral responses have not been previously considered as a mechanism of persistent hantavirus infections in reservoir hosts. This is due to observations of strong virus-specific IgG responses and high neutralizing antibody titers in hantavirus infected wild rodents (36–38). Therefore, our finding that PUUV-specific Ig responses were significantly delayed in voles infected with PUUV-Suo was unexpected, as was the lack of seroconversion in two of three bank voles infected with PUUV-wt. Interestingly, an acute increase in total IgG levels that cannot be solely explained by the PUUV-specific Ig response was observed for PUUV-Suo and PUUV-wt. An increase in total Ig levels, i.e. hypergammaglobulinemia, is strongly associated with other persistent viral infections in both humans and rodents (40–42). The mechanisms by which hypergammaglobulinemia facilitates persistent infections are currently unknown but could involve virus-mediated polyclonal activation of B cells and subsequent exhaustion of specific B cell responses (41). These could allow sufficient time for the virus to establish systemic infection that, despite later robust virus-specific humoral responses, the host is unable to clear.

Our results show that a hantavirus isolated using allogeneic reservoir host cells may retain wild-type characteristics better than a Vero E6-adapted virus, and should be considered as the preferred method for future hantavirus isolations and experimental infections. Using these techniques, we demonstrated that immune regulatory mechanisms are likely to facilitate persistent PUUV infections in bank voles, which provide insights into other difficult to study wildlife-hantavirus systems and may help guide novel therapeutic strategies for human infections.

## Materials and methods

### Ethics statement

All vole trapping and experimental procedures performed in Finland were approved the Animal Experimental Board of Finland (license number ESAVI/6935/04.10.07/2016). Wild bank voles representing the Eastern evolutionary lineage were captured in Ugglan live traps around central Finland. To generate the Mygla.REC.B cell line, an adult bank vole of the Western evolutionary lineage was taken from the breeding colony at the Friedrich-Loeffler-Institut, Greifswald-Insel Riems, Germany.

### Establishment of the permanent cell line

The bank vole from the breeding colony was anaesthetized with isoflurane and euthanized by using CO_2_. During dissection the kidney was removed and shipped in transport medium on ice to the Institute of Virology in Bonn, where the cell line was established. Briefly, the kidney was washed in sterile PBS and minced with a blade. Tissue was resolved with 37°C Dulbecco’s minimum essential medium-high glucose (DMEM; Sigma Aldrich) supplemented with 10 % inactivated fetal calf serum (FCS), 100 IU/ml Penicillin, 100 µg/ml Streptomycin, 2mM L-glutamine and a mix of non-essential amino acids (Sigma Aldrich), containing in addition 1% Ofloxain (Tarivid), and seeded into 6-well plates. After 5-7 days, primary cell growth could be observed, and medium was renewed. Primary cells were immortalized when confluency of 40-50% was reached by using a lentiviral system carrying the large T antigen of SV40, as described previously (43). Once an increase in cell proliferation was observed, cells were further passaged and cryoconserved.

### Virus isolation and propagation

For virus isolation, a serologically confirmed PUUV-infected bank vole from Suonenjoki, Finland and representing Eastern evolutionary lineage, was euthanized and lung samples were collected and frozen at −70 °C. The frozen lungs were then homogenized with a mortar and pestle in 1 ml of phosphate-buffered saline (PBS) over dry ice. After thawing, 500 µl of the homogenate was incubated for 1 h with semi-confluent Mygla.REC.B cells, which were grown similarly as described above. Mygla.REC.B cells were passaged at 3-day intervals until 100% of cells were found to be infected with PUUV via immunofluorescence assay as described below. We named the newly isolated hantavirus PUUV-Suonenjoki (PUUV-Suo) due to the geographical location of the bank vole trapping site.

An attenuated PUUV isolate, PUUV-Kazan strain (21), which was previously isolated and propagated in Vero E6 cells (green monkey kidney epithelial cell line; ATCC no. CRL-1586), served as comparison for experimental PUUV-Suo infections. PUUV-Kazan was grown in Vero E6 cells using Minimum Essential Medium (MEM; Sigma Aldrich) and supplemented with 10 % inactivated FCS, 100 IU/ml of Penicillin and 100 µg/ml of Streptomycin and 2mM of L-glutamine. Both PUUV-Suo and PUUV-Kazan strains were purified from supernatants of infected cells through a 30 % sucrose cushion by ultracentrifugation and resuspended in corresponding growth medium. Virus titers were measured by incubating diluted virus stocks with Vero E6 cells for 24 h at 37 °C, followed by acetone fixation and staining with a rabbit polyclonal antibody specific for PUUV N protein and AlexaFluor488-conjugated secondary antibody. Fluorescent focus-forming units (FFFU)/ml were counted under a UV microscope (Zeiss). UV inactivation of virus preparations was carried out using a Stratalinker UV crosslinker (300,000 μJ/cm2).

### Evolutionary lineage determination of bank voles and cell line

The lineage determination was based on an established protocol using amplification of a partial cytochrome *b* gene segment and comparison to prototype sequences of the various bank vole lineages (28, 44)

### Next-generation sequencing

Prior to RNA extraction, the bank vole lung homogenate and Mygla.REC.B cell culture supernatant were treated with a cocktail of micrococcal nuclease (New England BioLabs) and benzonase (Millipore) for 1 h at +37°C. RNA was extracted with Trizol and ribosomal RNA was removed using a NEBNext rRNA depletion kit (New England BioLabs), according to manufacturer instructions. The sequencing library was prepared using a NEBNext Ultra II RNA library prep kit (New England BioLabs) and quantified using a NEBNext Library Quant kit for Illumina (New England BioLabs). Pooled libraries were then sequenced on a MiSeq platform (Illumina) using a MiSeq v2 reagent kit with 150 bp paired-end reads. Raw sequence reads were trimmed and low-quality (quality score <30) and short (<50 nt) sequences were removed using Trimmomatic (45). Thereafter, *de novo* assembly was conducted using MegaHit, followed by re-assembly against the *de-novo* assembled consensus sequences using BWA-MEM implemented in SAMTools version 1.8 (46).

### Phylogenetic analysis

The complete L-, M- and S-segment sequences were downloaded from GenBank and aligned using MUSCLE program package (47). Substitution model was estimated using jModeltest2 (48). Phylogenetic trees were then constructed using the Bayesian Markov chain Monte Carlo (MCMC) method, implemented in Mr Bayes version 3.2 (49) using a GTR-G-I model of substitution with 2 independent runs and 4 chains per run. The analysis was run for 500 000 states and sampled every 5,000 steps.

### Cell cultures

PUUV-infected cells, grown on a black 96-well plate (Nunc), were detected by immunofluorescence assay after acetone fixation using a polyclonal rabbit anti-PUUN serum diluted 1:1000 in PBS followed by AlexaFluor488-conjugated donkey anti-rabbit secondary antibody (Thermo Scientific). Nuclei were stained using Hoechst 33420 diluted 1:5000 in PBS. Images were taken using a Leica TCS SP8 X confocal microscope (Biomedicum Imaging Unit core facility, Medicum, University of Helsinki).

Non- and PUUV-Suo infected Mygla.REC.B cells were exogenously stimulated with TLR-3 ligand polyI:C (10 µg/ml, Sigma Aldrich) or Sendai virus (multiplicity of infection 1; received from Prof. Ilkka Julkunen; National Institute of Health and Welfare, Helsinki, Finland). Cells were collected at 1 and 3-d post-treatment and RNA expression analysis was performed as described below. All experiments were conducted in duplicate.

### Experimental infections

Wild-captured voles from PUUV endemic region in Central Finland were transported to the University of Helsinki BSL-3 facility and acclimatized to individually ventilated biocontainment cages (ISOcage, Scanbur) for two days with *ad libitum* water and food (rodent pellets and small pieces of turnip every second day). Prior to any treatments, a small blood sample was collected from the retro-orbital sinus of each animal, and their PUUV infection status was determined by immunofluorescence assay (see below).

PUUV seronegative voles were divided into five treatment groups and subcutaneously injected with: 1) 100 µl of PBS (mock infection; n = 2 per time point), 2) 10,000 FFFUs of PUUV-Suo (n = 4 per time point), 3) UV-inactivated PUUV-Suo (n = 2 per time point), 4) 10,000 FFFUs of PUUV-Kazan (n = 2-3 per time point), or 5) a pooled homogenate prepared from the lungs of five PUUV-seropositive wild bank voles (PUUV-wt; n = 4 per time point). A total of 43 individual voles were used across the different time points and experimental treatments. The lung homogenate was obtained after grinding frozen lung tissue from naturally PUUV-infected voles (confirmed by PUUV-specific RT-PCR) with mortar and pestle in 1 ml PBS on dry ice. The PUUV-Suo and PUUV-wt were derived from voles captured from the same geographical area in Central Finland and together with voles used for experimental infections represent the Eastern bank vole evolutionary lineage. Blood samples from all treatment groups were collected from the retro-orbital sinus at 1wk intervals pi. Urine samples were collected from PUUV-Suo infected voles at 3d, and 1 and 2wks pi. Voles were sacrificed using isoflurane anesthesia, followed by cervical dislocation at 3d pi and 1 (7d), 2 (14-16d), and 5wks (35-38d) pi to collect samples for viral RNA load and distribution analyses, immunohistology and gene expression assays.

### Virus quantification

Following euthanasia and dissections, RNA extractions on bank vole tissues and urine were performed using Trisure (Bioline) according to the manufacturers’ instructions, with 10 µg/ml glycogen as carrier. RNA was directly subjected to PUUV S-segment RT-qPCR analysis based on a previously described protocol (50), with TaqMan fast virus 1-step master mix (Thermo scientific) using AriaMx instrumentation (Agilent).

### Histological and immunohistochemistry examinations

Two wild-trapped, naturally PUUV-infected adult bank voles were dissected and samples from brain, heart, lung, liver, kidneys and spleen were fixed in 10% neutral-buffered formalin. Similarly, lung, liver, spleen and kidney samples were collected and formalin-fixed from each two PUUV-Suo infected voles euthanized at 3d, 1wk, 2wks, and 5wks pi and two PUUV-wt infected bank voles euthanized at 5wks pi. The latter two voles were initially frozen at −80 °C, and tissue fixation was achieved by slowly thawing the organ samples in ice-cold formalin. After 4-7 days in formalin, tissue specimens were transferred to 70% ethanol, trimmed and routinely paraffin wax embedded. Consecutive sections (3-5 µm) were prepared and routinely stained with hematoxylin-eosin (HE) or subjected to immunohistology for the detection of PUUV N antigen in tissues.

Anti-PUUV N protein antiserum was generated by immunization of a single rabbit with PUUV N protein produced via baculovirus-mediated expression. The same batch of PUUV N protein was used in an earlier diagnostic study (51) and the immunization by BioGenes GmbH (Berlin, Germany) was as described in (52). Immunohistology was performed in an autostainer (Dako) using the custom-made rabbit polyclonal antiserum and the horseradish peroxidase (HRP) method. Briefly, sections were deparaffinized and rehydrated through graded alcohol. Antigen retrieval was achieved by 20 min incubation in citrate buffer (pH 6.0) at 98°C in a pressure cooker. This was followed by incubation with the primary antibody (diluted 1:1,000 in dilution buffer; Dako) for 60 min at room temperature (RT), a 10 min incubation at room temperature (RT) with peroxidase blocking buffer (Dako) and a 30 min incubation at RT with Envision+System HRP Rabbit (Dako). The reaction was visualized with diaminobenzidin (DAB; Dako). After counterstaining with hematoxylin for 2 sec, sections were dehydrated and placed on a coverslip with Tissue-Tek Film (Sysmex). A formalin-fixed and paraffin embedded pellet of Vero E6 cells infected with PUUV for 14 days served as a positive control (infected cells exhibit a finely granular to coarse cytoplasmic staining).

### PUUV-specific and total IgG assays

Immunofluorescence assays, using PUUV Sotkamo strain-infected Vero E6 cells fixed to microscope slides with acetone, were used to evaluate PUUV-specific Ig in bank vole blood (53). After incubating slides with blood diluted in PBS (1:10), bound Igs were detected with Fluorescein isothiocyanate (FITC)-conjugated rabbit anti-mouse Ig antibody (Dako Cytomation). To quantify total IgG levels, an enzyme-linked immunosorbent assay kit detecting mouse IgG (Mabtech) was used, replacing the mouse IgG standard with bank vole IgG 4G2 (specific for the PUUV glycoprotein) (54). One PUUV-wt infected vole had high total IgG levels already at the beginning of the experiment and was removed from the analysis.

### Splenocyte stimulations

To directly assess the effect of PUUV infection on immune cells, we extracted single cell suspensions from the homogenized spleens of 3 wild-caught PUUV-seronegative bank voles by washing the homogenate through 70 µM cell strainers (Sigma Aldrich) with 10 ml PBS. After pelleting through centrifugation, cells were incubated in 3 ml of Ammonium-Chloride-Potassium (ACK) lysing solution to lyse the red blood cells, washed in PBS and frozen in CryoStor CS10 (Sigma Aldrich) at −130°C until further investigation. After thawing and washing in PBS, splenocytes were suspended in RPMI-1640 supplemented with 10% inactivated FCS, 100 IU/ml of Penicillin, 100 µg/ml of Streptomycin, 2mM of L-glutamine and 25 mM Hepes pH 7.4. For stimulations, 1 million cells were either: 1) untreated, 2) treated with 20 µg/ml lipopolysaccharide (LPS), or 3) infected with virus at a m.o.i. of 0.01 in 250 µl of medium. Cells were collected after 3 days of incubation at 37 °C and subjected to RT-qPCR as described below. As an indicator of splenocyte viability after thawing, we used only splenocytes that showed a minimum of 5-fold increase in IFN-γ mRNA levels in response to LPS when compared to untreated, non-infected controls.

### Transcription factor and cytokine assays

To analyze bank vole gene expression profiles, RNA extracted from spleens and cell cultures was reverse transcribed to complementary DNA (cDNA) using random hexamers and RevertAid reverse transciptase (Thermo Scientific). Relative quantitative PCR was performed with Maxima SYBR Green master mix (Thermo Scientific) using AriaMx instrumentation (Agilent).

Primers used to measure the mRNAs levels of the interferon-inducible gene Mx2, a marker of innate immunity activation towards virus infection (55), and actin, used for normalization of mRNA levels between samples, are described in (56). Primers used to amplify bank vole mRNAs of Tbet, Gata3, FoxP3, IFN-γ, TGF-β, and IL-10 were originally designed for field voles and are described in (57). Primers to detect IL-4 mRNA (Forward: 5’-GCT CTG CCT TCT AGC ATG TAC T-3’ and Reverse: 5’-TGC ATG GCG TCC CTT TTT CT-3’) were designed based on the annotated mRNA sequence of the prairie vole (*Microtus ochrogaster*). All primers were validated for the corresponding bank vole sequence based on the expected amplicon sizes in agarose gels. Samples were excluded from further analysis when the PCR did not produce the expected single peak melting curve. Fold changes of individual mRNA expression levels relative to mock-infected controls were performed by the comparative CT method (58).

### Statistical analyses

Statistical differences between groups were assessed by one- or two-way ANOVA with Dunnett’s or Sivak’s multiple comparison tests as indicated in the figure legends. Analyses were conducted using GraphPad Prism version 8.1.2.

## Acknowledgements

We thank Sanna Mäki, Irina Suomalainen (University of Helsinki) and laboratory technicians in the Histology Laboratory at the University of Zurich for technical assistance. The authors would like to thank Markus Keller and Susanne Röhrs for provision of the bank vole, Matthias Lenk for dissection and provision of the kidney tissue for establishment of the Mygla.REC.B cell line, Dörte Kaufmann, Florian Binder and Stephan Drewes for determination of the evolutionary lineage by cytochrome *b* gene sequencing, and Stephan Drewes for helpful comments. This work was financially supported by the Academy of Finland (Grant 1275597 to TS, 1308613 to JH) and the Finnish Cultural Foundation (grant to KMF). The work of RGU (RoBoPub consortium) was supported by the Federal Ministry of Education and Research (BMBF) within the Research Network Zoonotic Infectious Diseases (FKZ 01KI1721A RGU). The work of IE (EpiZell) was funded by Federal Ministry of Education and Research (BMBF) through the National Research Platform for Zoonosis Research (FKZ 01KI1308).

## Supplementary figures

Supplementary Figure 1. Phylogenetic analysis of the complete PUUV-Suo S segment. Analysis of both the original bank vole lung (lung homogenate) and Mygla.REC.B.-isolated (cell culture supernatant) are included and marked as red.

Supplementary Figure 2. Phylogenetic analysis of the complete PUUV-Suo M segment. Analysis of both the original bank vole lung (lung homogenate) and Mygla.REC.B.-isolated (cell culture supernatant) are included and marked as red.

Supplementary Figure 3. Phylogenetic analysis of the complete PUUV-Suo L segment. Analysis of both the original bank vole lung (lung homogenate) and Mygla.REC.B.-isolated (cell culture supernatant) are included and marked as red.

## References

1. Plowright RK, Parrish CR, McCallum H, Hudson PJ, Ko AI, Graham AL, et al. Pathways to zoonotic spillover. Nat Rev Microbiol. 2017 Aug;15(8):502–10.

2. Viana M, Mancy R, Biek R, Cleaveland S, Cross PC, Lloyd-Smith JO, et al. Assembling evidence for identifying reservoirs of infection. Trends Ecol Evol. 2014 May;29(5):270–9.

3. Han BA, Park AW, Jolles AE, Altizer S. Infectious disease transmission and behavioural allometry in wild mammals. J Anim Ecol. 2015 May;84(3):637–46.

4. Plowright RK, Field HE, Smith C, Divljan A, Palmer C, Tabor G, et al. Reproduction and nutritional stress are risk factors for Hendra virus infection in little red flying foxes (Pteropus scapulatus). Proc Biol Sci. 2008 Apr 7;275(1636):861–9.

5. Altizer S, Bartel R, Han BA. Animal migration and infectious disease risk. Science. 2011 Jan 21;331(6015):296–302.

6. Forbes KM, Sironen T, Plyusnin A. Hantavirus maintenance and transmission in reservoir host populations. Curr Opin Virol. 2018 Feb;28:1–6.

7. Jonsson CB, Figueiredo LT, Vapalahti O. A global perspective on hantavirus ecology, epidemiology, and disease. Clin Microbiol Rev. 2010 Apr;23(2):412–41.

8. Vaheri A, Strandin T, Hepojoki J, Sironen T, Henttonen H, Makela S, et al. Uncovering the mysteries of hantavirus infections. Nat Rev Microbiol. 2013 Aug;11(8):539–50.

9. Vapalahti O, Mustonen J, Lundkvist A, Henttonen H, Plyusnin A, Vaheri A. Hantavirus infections in Europe. Lancet Infect Dis. 2003 Oct;3(10):653–61.

10. Kotlik P, Deffontaine V, Mascheretti S, Zima J, Michaux JR, Searle JB. A northern glacial refugium for bank voles (Clethrionomys glareolus). Proc Natl Acad Sci U S A. 2006 Oct 3;103(40):14860–4.

11. Filipi K, Markova S, Searle JB, Kotlik P. Mitogenomic phylogenetics of the bank vole Clethrionomys glareolus, a model system for studying end-glacial colonization of Europe. Mol Phylogenet Evol. 2015 Jan;82 Pt A:245–57.

12. Wójcik JM, Kawałko A, Marková S, Searle JB, Kotlík P. Phylogeographic signatures of northward post-glacial colonization from high-latitude refugia: a case study of bank voles using museum specimens. Journal of Zoology. 2010;281:249–62.

13. Deffontaine V, Libois R, Kotlik P, Sommer R, Nieberding C, Paradis E, et al. Beyond the Mediterranean peninsulas: evidence of central European glacial refugia for a temperate forest mammal species, the bank vole (Clethrionomys glareolus). Mol Ecol. 2005 May;14(6):1727–39.

14. Oldstone MB. Viral persistence: parameters, mechanisms and future predictions. Virology. 2006 Jan 5;344(1):111–8.

15. Easterbrook JD, Klein SL. Immunological mechanisms mediating hantavirus persistence in rodent reservoirs. PLoS Pathog. 2008 Nov;4(11):e1000172.

16. Schountz T, Prescott J. Hantavirus immunology of rodent reservoirs: current status and future directions. Viruses. 2014 Mar 14;6(3):1317–35.

17. Yanagihara R, Amyx HL, Gajdusek DC. Experimental infection with Puumala virus, the etiologic agent of nephropathia epidemica, in bank voles (Clethrionomys glareolus). J Virol. 1985 Jul;55(1):34–8.

18. Botten J, Mirowsky K, Kusewitt D, Bharadwaj M, Yee J, Ricci R, et al. Experimental infection model for Sin Nombre hantavirus in the deer mouse (Peromyscus maniculatus). Proc Natl Acad Sci U S A. 2000 Sep 12;97(19):10578–83.

19. Prescott J, Feldmann H, Safronetz D. Amending Koch’s postulates for viral disease: When “growth in pure culture” leads to a loss of virulence. Antiviral Res. 2017 Jan;137:1–5.

20. Safronetz D, Prescott J, Feldmann F, Haddock E, Rosenke R, Okumura A, et al. Pathophysiology of hantavirus pulmonary syndrome in rhesus macaques. Proc Natl Acad Sci U S A. 2014 May 13;111(19):7114–9.

21. Lundkvist A, Cheng Y, Sjolander KB, Niklasson B, Vaheri A, Plyusnin A. Cell culture adaptation of Puumala hantavirus changes the infectivity for its natural reservoir, Clethrionomys glareolus, and leads to accumulation of mutants with altered genomic RNA S segment. J Virol. 1997 Dec;71(12):9515–23.

22. Klingstrom J, Plyusnin A, Vaheri A, Lundkvist A. Wild-type Puumala hantavirus infection induces cytokines, C-reactive protein, creatinine, and nitric oxide in cynomolgus macaques. J Virol. 2002 Jan;76(1):444–9.

23. Plyusnina A, Razzauti M, Sironen T, Niemimaa J, Vapalahti O, Vaheri A, et al. Analysis of complete Puumala virus genome, Finland. Emerg Infect Dis. 2012 Dec;18(12):2070–2.

24. Razzauti M, Plyusnina A, Henttonen H, Plyusnin A. Microevolution of Puumala hantavirus during a complete population cycle of its host, the bank vole (Myodes glareolus). PLoS One. 2013 May 22;8(5):e64447.

25. Emeny JM, Morgan MJ. Regulation of the interferon system: evidence that Vero cells have a genetic defect in interferon production. J Gen Virol. 1979 Apr;43(1):247–52.

26. Nemirov K, Lundkvist A, Vaheri A, Plyusnin A. Adaptation of Puumala hantavirus to cell culture is associated with point mutations in the coding region of the L segment and in the noncoding regions of the S segment. J Virol. 2003 Aug;77(16):8793–800.

27. Boratynski Z, Melo-Ferreira J, Alves PC, Berto S, Koskela E, Pentikainen OT, et al. Molecular and ecological signs of mitochondrial adaptation: consequences for introgression? Heredity (Edinb). 2014 Oct;113(4):277–86.

28. Drewes S, Ali HS, Saxenhofer M, Rosenfeld UM, Binder F, Cuypers F, et al. Host-Associated Absence of Human Puumala Virus Infections in Northern and Eastern Germany. Emerg Infect Dis. 2017 Jan;23(1):83–6.

29. Binder F, Lenk M, Weber S, Stoek F, Dill V, Reiche S, et al. Common vole (Microtus arvalis) and bank vole (Myodes glareolus) derived permanent cell lines differ in their susceptibility and replication kinetics of animal and zoonotic viruses. J Virol Methods. 2019 Sep 9:113729.

30. Green W, Feddersen R, Yousef O, Behr M, Smith K, Nestler J, et al. Tissue distribution of hantavirus antigen in naturally infected humans and deer mice. J Infect Dis. 1998 Jun;177(6):1696–700.

31. Schountz T, Prescott J, Cogswell AC, Oko L, Mirowsky-Garcia K, Galvez AP, et al. Regulatory T cell-like responses in deer mice persistently infected with Sin Nombre virus. Proc Natl Acad Sci U S A. 2007 Sep 25;104(39):15496–501.

32. Easterbrook JD, Zink MC, Klein SL. Regulatory T cells enhance persistence of the zoonotic pathogen Seoul virus in its reservoir host. Proc Natl Acad Sci U S A. 2007 Sep 25;104(39):15502–7.

33. Rojas JM, Avia M, Martin V, Sevilla N. IL-10: A Multifunctional Cytokine in Viral Infections. J Immunol Res. 2017;2017:6104054.

34. Blackburn SD, Wherry EJ. IL-10, T cell exhaustion and viral persistence. Trends Microbiol. 2007 Apr;15(4):143–6.

35. Ejrnaes M, Filippi CM, Martinic MM, Ling EM, Togher LM, Crotty S, et al. Resolution of a chronic viral infection after interleukin-10 receptor blockade. J Exp Med. 2006 Oct 30;203(11):2461–72.

36. Brooks DG, Trifilo MJ, Edelmann KH, Teyton L, McGavern DB, Oldstone MB. Interleukin-10 determines viral clearance or persistence in vivo. Nat Med. 2006 Nov;12(11):1301–9.

37. Tian Y, Mollo SB, Harrington LE, Zajac AJ. IL-10 Regulates Memory T Cell Development and the Balance between Th1 and Follicular Th Cell Responses during an Acute Viral Infection. J Immunol. 2016 Aug 15;197(4):1308–21.

38. Snell LM, Osokine I, Yamada DH, De la Fuente JR, Elsaesser HJ, Brooks DG. Overcoming CD4 Th1 Cell Fate Restrictions to Sustain Antiviral CD8 T Cells and Control Persistent Virus Infection. Cell Rep. 2016 Sep 20;16(12):3286–96.

39. Marsac D, Garcia S, Fournet A, Aguirre A, Pino K, Ferres M, et al. Infection of human monocyte-derived dendritic cells by ANDES Hantavirus enhances pro-inflammatory state, the secretion of active MMP-9 and indirectly enhances endothelial permeability. Virol J. 2011 May 13;8:223,422X-8-223.

40. De Milito A, Nilsson A, Titanji K, Thorstensson R, Reizenstein E, Narita M, et al. Mechanisms of hypergammaglobulinemia and impaired antigen-specific humoral immunity in HIV-1 infection. Blood. 2004 Mar 15;103(6):2180–6.

41. Hunziker L, Recher M, Macpherson AJ, Ciurea A, Freigang S, Hengartner H, et al. Hypergammaglobulinemia and autoantibody induction mechanisms in viral infections. Nat Immunol. 2003 Apr;4(4):343–9.

42. Kawamoto H, Sakaguchi K, Takaki A, Ogawa S, Tsuji T. Autoimmune responses as assessed by hypergammaglobulinemia and the presence of autoantibodies in patients with chronic hepatitis C. Acta Med Okayama. 1993 Oct;47(5):305–10.

43. Eckerle I, Ehlen L, Kallies R, Wollny R, Corman VM, Cottontail VM, et al. Bat airway epithelial cells: a novel tool for the study of zoonotic viruses. PLoS One. 2014 Jan 13;9(1):e84679.

44. Schlegel M, Ali HS, Stieger N, Groschup MH, Wolf R, Ulrich RG. Molecular identification of small mammal species using novel cytochrome B gene-derived degenerated primers. Biochem Genet. 2012 Jun;50(5-6):440–7.

45. Bolger AM, Lohse M, Usadel B. Trimmomatic: a flexible trimmer for Illumina sequence data. Bioinformatics. 2014 Aug 1;30(15):2114–20.

46. Li H, Handsaker B, Wysoker A, Fennell T, Ruan J, Homer N, et al. The Sequence Alignment/Map format and SAMtools. Bioinformatics. 2009 Aug 15;25(16):2078–9.

47. Edgar RC. MUSCLE: a multiple sequence alignment method with reduced time and space complexity. BMC Bioinformatics. 2004 Aug 19;5:113,2105-5-113.

48. Darriba D, Taboada GL, Doallo R, Posada D. jModelTest 2: more models, new heuristics and parallel computing. Nat Methods. 2012 Jul 30;9(8):772.

49. Ronquist F, Teslenko M, van der Mark P, Ayres DL, Darling A, Hohna S, et al. MrBayes 3.2: efficient Bayesian phylogenetic inference and model choice across a large model space. Syst Biol. 2012 May;61(3):539–42.

50. Niskanen S, Jaaskelainen A, Vapalahti O, Sironen T. Evaluation of Real-Time RT-PCR for Diagnostic Use in Detection of Puumala Virus. Viruses. 2019 Jul 19;11(7):10.3390/v11070661.

51. Hepojoki S, Hepojoki J, Hedman K, Vapalahti O, Vaheri A. Rapid homogeneous immunoassay based on time-resolved Forster resonance energy transfer for serodiagnosis of acute hantavirus infection. J Clin Microbiol. 2015 Feb;53(2):636–40.

52. Korzyukov Y, Hetzel U, Kipar A, Vapalahti O, Hepojoki J. Generation of Anti-Boa Immunoglobulin Antibodies for Serodiagnostic Applications, and Their Use to Detect Anti-Reptarenavirus Antibodies in Boa Constrictor. PLoS One. 2016 Jun 29;11(6):e0158417.

53. Kallio-Kokko H, Laakkonen J, Rizzoli A, Tagliapietra V, Cattadori I, Perkins SE, et al. Hantavirus and arenavirus antibody prevalence in rodents and humans in Trentino, Northern Italy. Epidemiol Infect. 2006 Aug;134(4):830–6.

54. Lundkvist A, Niklasson B. Bank vole monoclonal antibodies against Puumala virus envelope glycoproteins: identification of epitopes involved in neutralization. Arch Virol. 1992;126(1-4):93–105.

55. Haller O, Kochs G. Human MxA protein: an interferon-induced dynamin-like GTPase with broad antiviral activity. J Interferon Cytokine Res. 2011 Jan;31(1):79–87.

56. Stoltz M, Sundstrom KB, Hidmark A, Tolf C, Vene S, Ahlm C, et al. A model system for in vitro studies of bank vole borne viruses. PLoS One. 2011;6(12):e28992.

57. Jackson JA, Begon M, Birtles R, Paterson S, Friberg IM, Hall A, et al. The analysis of immunological profiles in wild animals: a case study on immunodynamics in the field vole, Microtus agrestis. Mol Ecol. 2011 Mar;20(5):893–909.

58. Schmittgen TD, Livak KJ. Analyzing real-time PCR data by the comparative C(T) method. Nat Protoc. 2008;3(6):1101–8.

